# Motor and Visual Plasticity interact in Adult Humans

**DOI:** 10.1101/2022.05.03.490377

**Authors:** Izel D. Sarı, Claudia Lunghi

**Affiliations:** Laboratoire des systèmes perceptifs, Département d’études cognitives, École normale supérieure, PSL University, CNRS, 75005 Paris, France

## Abstract

Neuroplasticity is maximal during development and declines in adulthood, especially for sensory cortices. On the contrary, the motor cortex retains plasticity throughout the lifespan. This difference has led to a modular view of plasticity in which different brain regions have their own plasticity mechanisms that do not depend or translate on others. Recent evidence indicates that visual and motor plasticity share common neural mechanisms (e.g. GABAergic inhibition), indicating a possible link between these different forms of plasticity, however the interaction between visual and motor plasticity has never been tested directly. Here we show that when visual and motor plasticity are elicited at the same time in adult humans, visual plasticity is impaired, while motor plasticity is spared. This unilateral interaction between visual and motor plasticity demonstrates a clear link between these two forms of plasticity. We conclude that local neuroplasticity in separate systems might be regulated globally, to preserve overall homeostasis in the brain.

## Introduction

Neuroplasticity in the sensory brain is at its peak during the critical period in development and steeply decreases afterwards^1, 2^. The visual cortex is one of the most widely used models to study neuroplasticity, with the gold standard paradigm of ocular dominance (OD) plasticity: during the critical period, monocular deprivation (MD) shifts neural responses in the primary visual cortex in favor of the open eye and leaves the deprived eye amblyopic^3–8^. After the closure of the critical period, this OD change is no longer observed and the visual cortex is thought to become hardwired^3, 9^. On the other hand, the adult motor cortex retains a high level of plasticity throughout the lifespan, as seen for example in motor learning^10^. This disparity in the plastic potentials of different brain regions in adulthood led to a modular approach to the study of neuroplasticity assuming different brain regions have their own plasticity mechanisms that do not depend or translate on others.

Growing evidence is showing that the adult human visual cortex retains a particular form of OD plasticity: after a few hours of monocular deprivation, the deprived eye unexpectedly increases its dominance over the non-deprived eye, as measured by binocular rivalry^11–14^ and other tasks^15–18^. This counterintuitive OD shift also occurs at the neural level: short-term MD boosts deprived-eye related activity in the early visual cortex, while reducing responses to the non-deprived eye^13, 15, 19–22^. Moreover, this effect is accompanied by a decrease in resting GABA concentration in the visual cortex^14^, suggesting the involvement of a homeostatic mechanism that aims preservation of a constant Excitation/Inhibition (E/I) balance^23^. Interestingly, motor learning is also closely tied to GABA modulation in the motor cortex^24^. Decreased GABA concentration in the motor and sensorimotor cortices facilitates motor learning^25, 26^ and individual variations in motor learning performance correlate with the changes in motor cortex GABA concentration following transcranial direct current stimulation^27^.

That the adult visual system retains a degree of homeostatic plasticity higher than previously thought and that visual and motor plasticity share common neural mechanisms (E/I balance alteration) suggest that these forms of plasticity might be linked and controlled jointly to ensure overall stability in the brain. However, this hypothesis has never been tested directly. Here we address this issue by investigating the interaction between visual and motor plasticity in adult humans using two well-established experimental paradigms to elicit each form of plasticity (i.e. short-term monocular deprivation^11, 15^ and motor sequence learning^10, 28,29,30,31^). We designed a combined task in which visual and motor plasticity are induced at the same time and compared visual and motor plasticity in this condition to simple tasks in which either form of plasticity was induced on its own. Our results show a unilateral interaction between visual and motor plasticity that impairs visual plasticity in the presence of active motor plasticity.

## Materials & Methods

### Subjects

34 participants (Mean age 28.3 ± 4.8 years, 12 Males) with normal or corrected-to normal visual acuity (measured with ETDRS charts) took part in the study. One subject was excluded from the analysis due to inconsistent ODI within and across the binocular rivalry blocks on the same day, two others were excluded due to high variation in baseline ocular dominance on different days.

Data from the remaining 31 participants was analyzed. All subjects but two (the authors) were naïve to the purpose of the experiment.

### Ethics Statement

The experimental protocol was approved by the local ethics committee (Comité d’éthique de la Recherche de l’université Paris Descartes, CER-PD:2019-16-LUNGHI) and was performed in accordance with the Declaration of Helsinki (DoH-Oct2008). All participants gave written informed consent. The naïve participants received financial compensation of 10€ per hour.

### Apparatus and Stimuli

#### Visual task: binocular rivalry

The experiment took place in a dark and quiet room. The visual stimuli were generated in Matlab (R2020b, The MathWorks Inc., Natick, MA) using Psychtoolbox-3^32^ running on a PC (Alienware Aurora R8, Alienware Corporation, Miami, Florida, USA) and a NVIDIA graphics card (GeForce RTX2080, Nvidia Corporation, Santa Clara, California, USA).

Visual stimuli were presented dichoptically through a custom-built mirror stereoscope and each subject’s head was stabilized with a forehead and chin rest positioned 57 cm from the screen. Visual stimuli were two sinusoidal gratings oriented either 45° clockwise or counterclockwise (size: 3.1°, spatial frequency: 2 cpd, contrast: 50%), presented on a uniform gray background (luminance: 110 cd/m2, CIE x=.305, y=.332) in central vision with a central white fixation point and a common squared white frame to facilitate dichoptic fusion. The stimuli were displayed on an LCD monitor (BenQ XL2420Z 1920 x 1080 pixels, 144 Hz refresh rate, Tapei, Taiwan).

Responses were recorded through the computer keyboard. Monocular deprivation was performed using eye-patching. The eye-patch was made of a translucent plastic material that allowed light to reach the retina but prevented pattern vision, in accordance with previous studies^11, 13, 33^.

#### Motor task: motor sequence learning

The motor learning experiment took place in a quiet and lit room. The experiment ran on a PC (Dell) with Matlab using Psychtoolbox-3. A series of numbers in white (or lines of a story for the visual control condition) were presented on a uniform gray background (luminance: 47.6 cd/m2, CIE x=.306, y=.335). Observers viewed the display at a distance of 50 cm on an LCD monitor (Dell 2016H, 1600 x 900 pixels, 60 Hz refresh rate, Dell Inc., Round Rock, Texas, USA). Responses were recorded through a four-buttoned response pad (4-button USB response pad carbon fibre effect, The Blackbox Toolkit Ltd., Sheffield, UK).

#### Attentional control: simple visual and working memory tasks

The experiment ran on a PC (Dell OptiPlex 7070, Dell Inc., Round Rock, Texas, USA) equipped with NVIDIA graphics card (GeForce GTX 1050, Nvidia Corporation, Santa Clara, California, USA), and the stimuli were generated with Matlab using Psychtoolbox-3. We used an LCD monitor (ViewSonic V3D245 1920 x 1080 pixels, 120 Hz refresh rate, ViewSonic Corporation, Brea, California, USA) to display the stimuli. For the simple visual task, the stimuli were the same sinusoidal gratings described above. For the working memory task, participants saw a series of white letters briefly flashed on a uniform gray background (luminance: 44.8 cd/m2, CIE x=.299, y=.340). All participants observed the display from a 57cm distance, stabilized by a chin rest, in a quiet and dark room. Responses were recorded through the computer keyboard.

### Procedures

All participants performed three different conditions: simple motor, simple visual and combined. The order of conditions was counterbalanced across subjects and each condition was performed on separate days. Because of recent evidence showing an interaction between visual homeostatic plasticity and energy metabolism^34^, each condition was performed at approximately the same time of the day and immediately after lunch.

#### Simple Visual Task

Ocular dominance was measured by means of binocular rivalry^35, 36^ . Visual plasticity was quantified as the ocular dominance shift occurring for each participant after short-term (150 minutes) monocular deprivation (Figure 1a). Participants viewed a screen through a mirror stereoscope and reported the change in their perception via the keyboard by pressing one of three arrow keys continuously according to the orientation of the grating they perceived at the moment (left arrow: counter-clockwise, right arrow: clockwise, down arrow: mixed). Each binocular rivalry block lasted 8 minutes including two 3-minute trials with a pause in between. The 3-minute trials consisted of two sub-trials of 90 seconds and a 3-second break in the middle. Participants remained in position through a first trial, a 2-minute break and a second trial of rivalry. After each break, grating locations were switched to avoid adaptation. The starting locations of the gratings were counter-balanced across participants.

**Figure 1.**
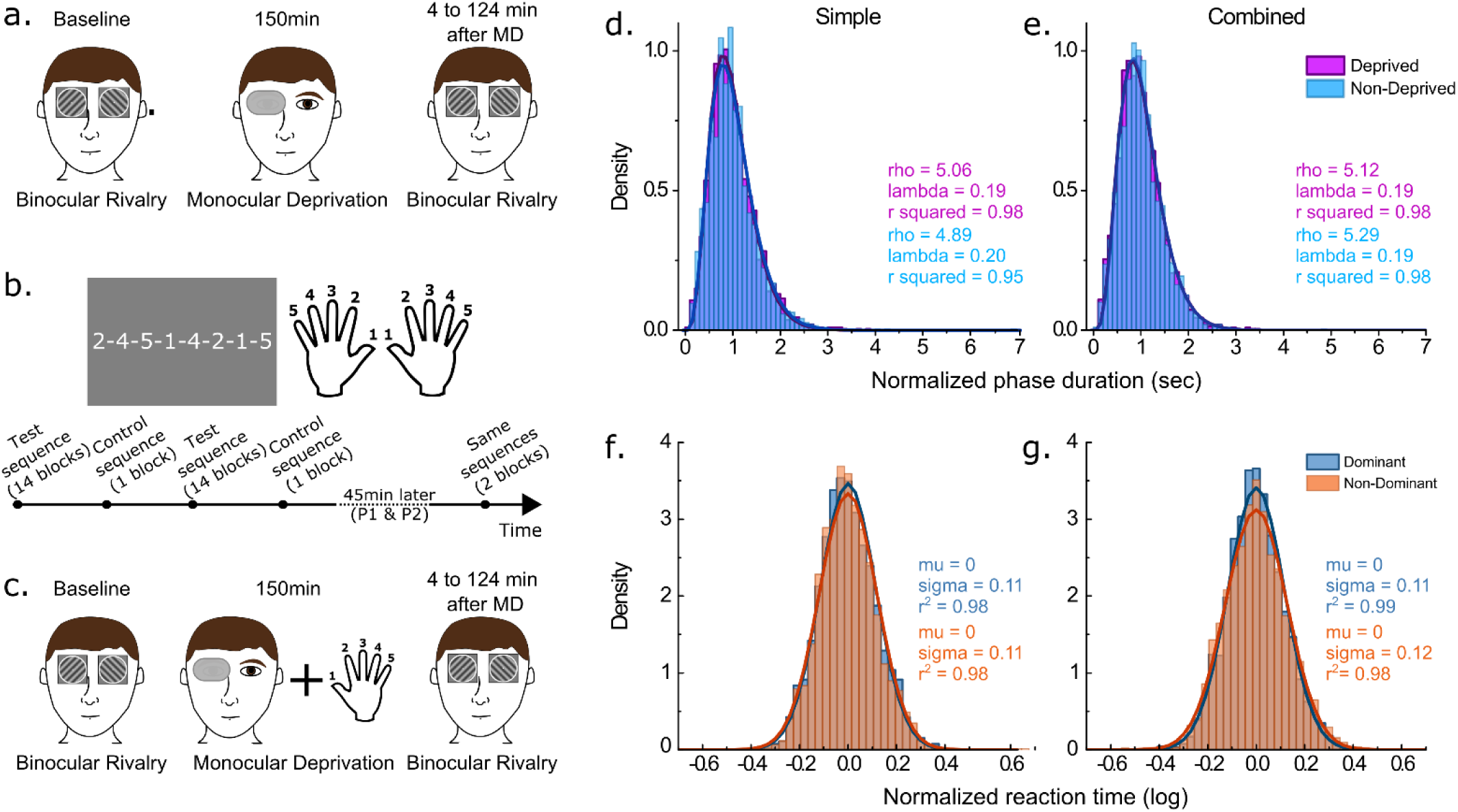
Experimental paradigm and response distributions. a) In the simple visual condition, participants performed a binocular rivalry task before and after 150 minutes of monocular deprivation. Participants indicated the orientation of the grating they perceive. b) In the simple motor task, participants were instructed to tap their fingers in the order displayed on the screen. Numbers corresponded to fingers. Each hand was trained separately. For each hand, participants practiced four different sequences: two test sequences (14 blocks) and two control sequences (1 block). 45 minutes after the initial training of a hand, participants performed the same four sequences for two blocks each. c) In the combined condition, participants performed the motor task during the monocular deprivation (MD) period. Binocular rivalry was performed before and after the MD period. d-e) Normalized phase duration distributions in de baseline blocks of the simple and combined visual conditions. The distributions are fitted with a gamma function (Eq.1). f-g) Log-transformed reaction time distributions, normalized across participants, in the first block of simple and combined motor conditions. The distributions are fitted with a Gaussian function (Eq.3).

After a baseline block of binocular rivalry, the participants wore an eye-patch for 150 minutes over their dominant eye. We used as operational definition of dominant eye the eye that dominated perception in binocular rivalry during the baseline block. When the ocular dominance in the baseline block was inconclusive, we used the PORTA test^37^ . During the 150 minutes of monocular deprivation, participants were free to perform different activities in the lab like work, read or browse the web or even go for a walk outside. At the end of this period the eye-patch was removed and the participants performed seven more blocks of binocular rivalry. The first five blocks were separated by 7 minutes while the last three blocks had 22 minutes in between. In total, ocular dominance was measured for 128 minutes following eye-patch removal. We calculated a unique ocular dominance index (ODI, see Eq 2) for each of the eight blocks. The ocular dominance index difference from the baseline at each time point served as a measure of visual plasticity.

#### Simple Motor Task

A motor sequence learning task was used to assess motor plasticity. Motor plasticity for each participant was quantified as the decrease in reaction time after 14 blocks of practice. Participants were instructed to tap their fingers in the order of the eight-element number sequence shown on the monitor. Each number on the screen corresponded to a finger and the numbering was mirrored, that is, the thumb was always number 1 regardless of the hand (Figure 1b). Our response pad having only four buttons, the number three was never shown on the screen so the middle finger was never used. Half of the participants started the training with their dominant hand and the other half with their non-dominant hand. Participants trained on four different finger tap sequences per hand (eight in total): two “test sequences” (repeated for 14 blocks) and two “control sequences” (repeated for only one block). The labeling and the order of these sequences were counterbalanced across participants.

The motor learning task consisted in a total of 30 blocks (14x2 test sequences and 1x2 control sequences) per hand. Each block consisted of five trials and each trial consisted of three sub-trials each corresponding to a full repetition of a finger tap sequence. Participants were instructed to tap their fingers in the displayed order, as rapidly and as correctly as possible. When the training of all four sequences were complete for a hand, participants took a break before starting the training of the other hand. About 45 minutes after the training of a hand, participants repeated all 4 sequences for 2 more blocks each to make sure the reaction time (RT) decrease was stable and specific to test sequences. The whole task lasted about 150 minutes.

Reaction times (time elapsed from the first until the last finger tap in a sub-trial) and accuracy were computed per each participant and for each experimental block. The reaction time decrease for test sequences from the first to the 14^th^ block served as a measure of motor plasticity.

#### Combined Task

In the combined condition, the simple visual and the motor tasks were performed at the same time (Figure 1c). Participants started with the baseline block of binocular rivalry. After the baseline block, participants wore the eye-patch. Here, instead of engaging in their preferred activity, all participants performed the motor learning task during the monocular deprivation period. This way, we induced motor and visual plasticity simultaneously. At the end of the monocular deprivation and the motor task, we removed the eye-patch and the participants completed 7 more binocular rivalry blocks. The same measures (RT, accuracy and ODI) were recorded.

#### Visual Control Task

In order to eliminate the possibility of a confound in our study, we included a fourth (control) condition. 10 participants from the main experiment cohort of observers (Mean age 27.3±3.6 years, 5M) took part in the visual control condition. In this control study, the effect of short-term monocular deprivation was measured as in the simple visual task. However, during the 150 minutes of monocular deprivation, participants read stories (white text presented on a uniform grey background), to match the visual stimulation received during the combined task. We used the same screen and background as in the motor task and the stories were displayed maximum two lines at a time (Figure 3a).

#### Attentional Control Task

To control for the possible effect of attentional and working memory load on monocular deprivation induced ocular dominant shift, we included another control condition in our study. 10 participants (mean age 28.3±5.7, 8F) participated in this attentional control condition. The effect of short-term monocular deprivation was measured as in the simple visual task. Participants performed a working memory task while they were under monocular deprivation (Figure 5a). A set of three, five or seven letters were flashed sequentially and briefly (0.15sec each) on a gray screen with an inter stimulus interval of 1.5 seconds. Participants had a 3 second memory retention period at the end of the set and before the flashing of the probe letter. The probe letter was equally likely to be within or outside of the set on each trial. Participants indicated whether the flashed probe letter belonged to the set (left arrow key) or not (right arrow key). The response was followed by an auditory feedback indicating correct (high pitch) or incorrect (low pitch) response. Each block consisted in three trials containing 30 sub-trials each. One block lasted about fifteen minutes. Participants started the first block of the working memory task at 15 minutes into the monocular deprivation period. In total, they completed 4 blocks of working memory task with 20 minute breaks in between. The starting condition (simple visual or attentional control) was counterbalanced across participants.

### Statistical Analyses

The recorded reaction times and perceptual durations were first analyzed on Matlab and then the statistical tests were performed with SPSS statistics (version 25, IBM Corporation, Armonk, New York, USA). The mean reaction times and ODI based on perceptual durations were separately compared with repeated measures ANOVAs. Although both the initial distribution of our data and the residuals of the ANOVAs significantly differed from normality, literature has shown that ANOVAs are quite robust in Type I & II errors and power with non-normal distributions ^38, 39^ and are arguably preferable to alternatives ^40^. Pairwise comparisons are made with Wilcoxon signed rank tests.

#### Visual Task

For the analysis of the binocular rivalry data, we discarded phase durations < 0.25sec to eliminate keypress errors. To plot perceptual phase duration distributions for the deprived and non-deprived eyes in the baseline blocks of the two different conditions, we first normalized each participant’s phase duration to their own mean. The distributions of the pooled data of all participants then were fitted by a two-parameter Gamma distribution (r, λ) of the form:

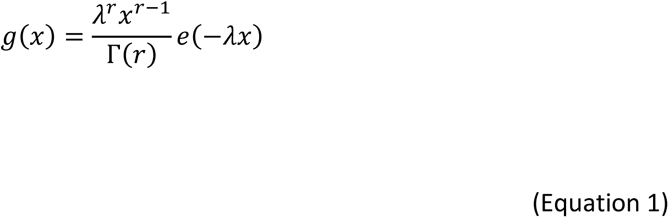

where Γ is the *gamma function*, *r* is the shape parameter and *λ* is the scale parameter. The goodness of fit (r-squared) was > 0.97 for all distributions (Figure 1d-e).

We compared the ocular dominance index change in the two conditions (simple vs combined). We calculated the ocular dominance index (ODI) as follows:

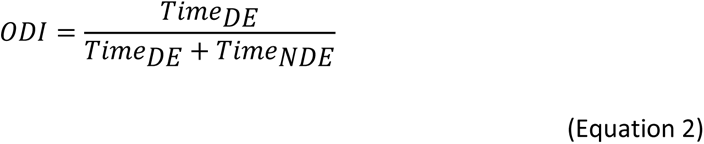

Where Time_DE_ indicates the total amount of time (in seconds) spent by the observer reporting the dominance of the stimulus presented to the deprived eye and Time_NDE_ indicates the total amount of time (in seconds) spent by the observer reporting the dominance of the stimulus presented to the non-deprived eye. ODI values around 0.5 indicate, therefore, balanced ocular dominance, while ODI > 0.5 indicates predominance of the deprived eye and ODI < 0.5 predominance of the non-deprived eye. We excluded the participants whose baseline ODI on the two days showed a difference larger than 0.12. The reason for this exclusion criterion is that the effect of monocular deprivation on the ocular dominance change is estimated to be about a 0.12 change in ODI based on the literature ^33^. Data from participants who show a natural variation larger than this value would not reflect the effect of monocular deprivation.

We subtracted the baseline ODI of each participant from the subsequent block ODIs to see the effect of patching on ocular dominance. We then compared the ODI change in two conditions (simple vs combined) in a 7 (Blocks) x 2 (Conditions) repeated-measures ANOVA. The binocular rivalry blocks taking place after the monocular deprivation period were binned: 0-8min, 15- 23min, 30-38min, 45-53min, 60-68min, 90-98min, 120-128min.

#### Motor Task

In the analysis of reaction times (RT), only the correct trials were taken into account (incorrect trials 13.7% for simple motor and 13.9% for combined condition). Outlier RTs (defined as the RTs µ ∓ 4sd of each participant) were removed (0.3% each for both simple motor and combined conditions). We looked at the reaction time distributions for dominant and non-dominant hands within the first block of each condition by first dividing each participant’s data to their mean RT. The distributions based on these raw reaction times showed very high skewness and kurtosis (Supplementary Figure 1). To work around this, the reaction times were log-transformed and normalized by subtracting the mean of each participant from their data. The pooled reaction time distributions were then fitted by a two-parameter Gaussian distribution (µ, σ) of the form:

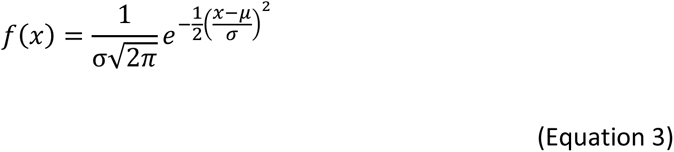

Where f(x) is the probability density function, µ is the mean and σ is the standard deviation. The goodness of fit (r-squared) was > 0.98 for all distributions (Figure 1f-g).

Working with the log-transformed reaction times, we calculated each participant’s mean reaction time in each block. We then subtracted each participant’s mean reaction time in the first block from all others, equalizing all participants to a starting point of 0.

We compared the mean reaction time differences from the 1^st^ to the 14^th^ block for only test sequences (dominant and non-dominant hand together) across the two conditions (simple vs combined) using a 13 (Blocks) x 2 (Conditions) repeated-measures ANOVA.

#### Correlation

To check for correlation between visual and motor plasticity, we used the data from simple visual and simple motor tasks. The visual plasticity score for each participant was calculated by subtracting the baseline ODI from the ODI immediately after monocular deprivation (0-8 min). The motor plasticity score for each participant was calculated by subtracting the mean RT difference from baseline on the 14^th^ or the 2^nd^ block from 0. Correlations between visual and motor plasticity scores were computed using the Spearman’s rank correlation coefficient (rho) and p-values by using the permutation distributions.

## Results

We assessed visual homeostatic plasticity and motor plasticity in a group (N=31) of adult volunteers using two well established experimental paradigms. We quantified visual homeostatic plasticity as the shift in ocular dominance (measured by binocular rivalry) occurring after 150 minutes of monocular deprivation (simple visual task, Figure 1a). The ocular dominance index (ODI, see Eq 2) measured before deprivation (baseline ODI) in the simple and combined conditions correlated across participants (rho = 0.36, p = 0.04, Supplementary Figure 2a), as well as mean phase durations (rho = 0.84, p < 0.0001, Supplementary Figure 2b) and mixed percept proportions (rho = 0.74, p < 0.0001, Supplementary Figure 2c), indicating that the ODI is a reliable measure of ocular dominance. We assessed motor plasticity as the reaction time decrease over fourteen blocks of finger-tap-sequence learning (simple motor task, Figure 1b). In a third condition, visual and motor plasticity were elicited at the same time, as participants performed the motor task during the 150 minutes of monocular deprivation (combined task, Figure 1c). Phase duration distributions in binocular rivalry and reaction time distributions for the motor learning task in the different experimental conditions are reported in Figure 1d-g. The phase duration distributions for binocular rivalry follow a gamma distribution (r-squared > 0.97) as expected^35^. The log-transformed reaction time distributions are well fit by normal distribution (r-squared > 0.98).

Results from the simple visual and motor task replicate the effects established in the literature^10, 11^. Ocular dominance shifts in favor of the deprived eye after monocular deprivation (Wilcoxon signed ranks z = 4.56, p < 0.001) and slowly returns to baseline after patch removal (Figure 2a, red symbols). Reaction times consistently decrease over fourteen blocks of learning for the motor task (Wilcoxon signed ranks z = -4.85, p < 0.001) and this decrease is maintained 45 minutes after initial training (Wilcoxon signed ranks z = -4.85, p < 0.001, Figure 2b, red symbols). The reaction times of test sequences remained significantly lower than those of control sequences 45 minutes after training (P1: Wilcoxon signed rank z = -3.1, p < 0.01, P2: Wilcoxon signed rank z = -3.7, p < 0.0001, Supplementary Figure 3).

**Figure 2.**
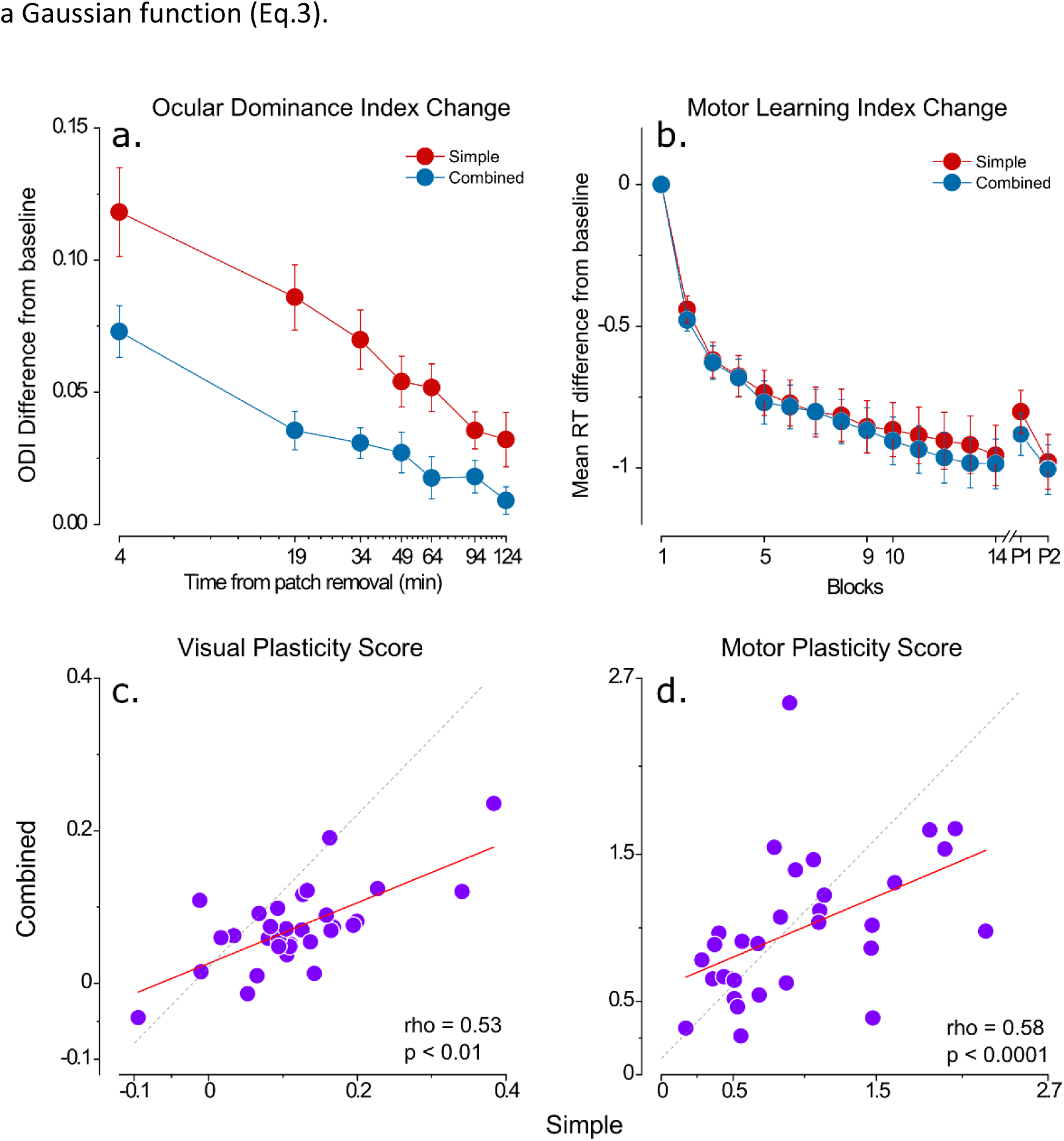
Results. a) Ocular dominance index (ODI) difference from baseline for the simple visual (red symbols) and combined (blue symbols) conditions. Error bars represent 1±S.E.M. b) Change in reaction times (RT) during motor learning (difference from baseline mean RT) in the simple (red symbols) and combined (blue symbols) conditions. Error bars represent 1±S.E.M. c) Scatter plot of the ODI difference measured during the first 8 minutes after monocular deprivation in the simple vs combined condition. d) Scatter plot of the RT change from baseline in 14^th^ Block in the simple vs combined condition.

Interestingly, in the combined condition, we observed a significantly smaller shift in ocular dominance following short-term monocular deprivation compared to the simple visual task (Figure 2a, blue symbols), indicating that eliciting visual and motor plasticity at the same time impaired the visual homeostatic response to monocular deprivation. A two-way (Block x Condition) repeated measures ANOVA showed a significant main effect of both the factor *Block* (F(6,180) = 27.9, p < .001, η^2^p = .48) and *Condition* (F(1,30) = 14.4, p = 0.001, η^2^p = 0.32) and a significant interaction between the two (F(6,180) = 2.76, p = 0.013, η^2^p = 0.08). The ODI change in the block immediately after monocular deprivation (i.e. visual plasticity score) was correlated across the two conditions (r = 0.68, p < 0.001) although the change in the simple condition was larger for most of the participants (Figure 2c), indicating a good test-retest reliability of binocular rivalry in quantifying the effect of short-term monocular deprivation. No significant difference in motor learning was observed in the combined condition compared to the simple task (Figure 2b, blue symbols): a two-way (Block x Condition) repeated measures ANOVA revealed a significant main effect of the factor *Block* (F(12,360) = 49.9, p < 0.0001, η^2^p = 0.62) whereas the factor *Condition* (F(1,30) = 0. 01, p > 0.05, η^2^p = 0.0) and the interaction of the two (F(12,360) = 0.44, p > 0.05, η^2^p = 0.01) remained insignificant. The mean reaction time difference from the baseline on the 14^th^ block (i.e. motor plasticity score) correlates across the two conditions (rho = 0.58, p < 0.0001, Figure 2d). In both conditions, participants performed the motor task with high accuracy rates throughout the training (mean = 0.87, sd = 0.10). We found no significant difference between the reaction times of the dominant and the non-dominant hands nor a difference between the first-trained vs second-trained hands or sequences. When visual and motor plasticity are elicited simultaneously, motor plasticity is not impacted.

Having shown that visual and motor plasticity interact with each other, we investigated whether a correlation exists between the plastic potentials of the visual and motor cortex. We found no correlation between visual and motor plasticity across participants. We calculated two plasticity scores, one representing the maximal effect of motor learning (difference of the mean reaction time on the last block from the first block of training) and one indicating the speed of learning (difference in mean reaction times from the first to the second block). Neither of these motor plasticity scores correlates with the visual plasticity score (r = -0.01, p > 0.05, Figure 3a, r= 0.01, p > 0.05, Figure 3b).

**Figure 3.**
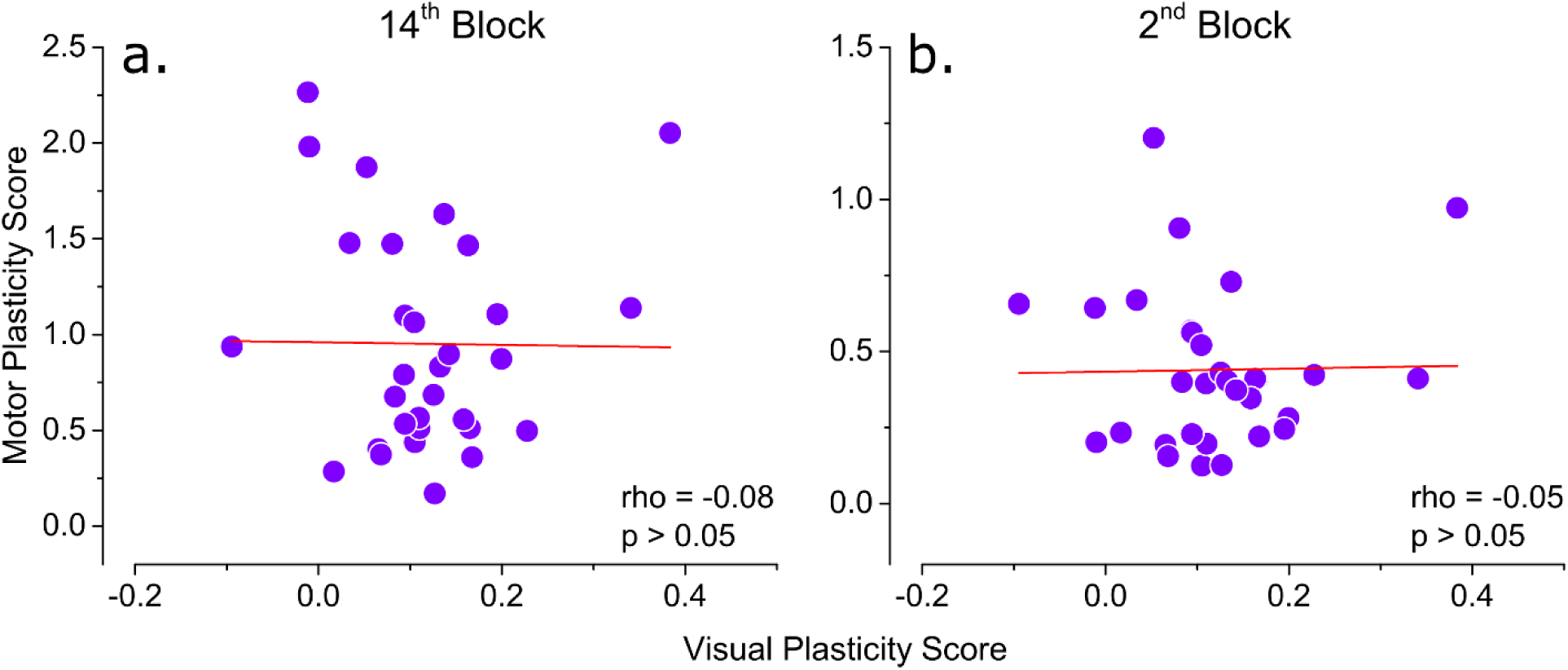
Correlation between visual and motor plasticity. a) Correlation between the visual plasticity score (ODI change in 8-minutes-after-deprivation) and motor plasticity score based on mean reaction time difference from baseline in the last block (0 – mean RT difference in 14^th^ block). b) Correlation between the visual plasticity score (same as panel A) and motor plasticity score based on reaction time difference from baseline in the second block (1 – mean RT difference in 2^nd^ block).

One possible confound, which could potentially affect the ocular dominance shift induced by monocular deprivation, is the difference in visual stimulation during monocular deprivation for the simple visual task and the combined task as shown by a previous study^34^. While in the simple visual task, during monocular deprivation participants were free to perform their preferred activities and had therefore access to a rich visual environment, in the combined condition, visual stimulation during deprivation was reduced to the finger-tap sequence numbers displayed on a grey background. In order to address this possible confound, we performed a control experiment in which we measured the effect of 150 minutes of monocular deprivation on 10 participants who also performed the main experiment. In this visual control condition, during monocular deprivation, participants read stories displayed on a grey background reproducing the same amount of visual stimulation as in the combined condition (Figure 4a). We found that the shift in ocular dominance observed in the control condition was comparable to that observed in the simple visual task (Figure 4b). A two-way ANOVA (Block x Condition) confirmed that while in both conditions the ocular dominance index changed with blocks (F(6,54) = 13.6, p < 0.001, η^2^p = 0.60), there was no significant effect of condition (F(1,9) = 13.6, p > 0.05, η^2^p = 0.60), nor interaction between the two (F(6,54) = 0.94, p > 0.05, η^2^p = 0.09). A two-way (Block x Condition) repeated measures ANOVA between control and combined conditions showed a significant main effect of both the factor *Block* (F(6,54) = 9.78, p < .001, η^2^p = .52) and *Condition* (F(1,9) = 9.55, p = 0.013, η^2^p = .51) while the interaction between the two (F(6,54) = .59, p > .05, η^2^p = .06) remained insignificant. This indicates that the drop in the monocular deprivation effect observed in the combined condition was not due to poor visual stimulation.

**Figure 4.**
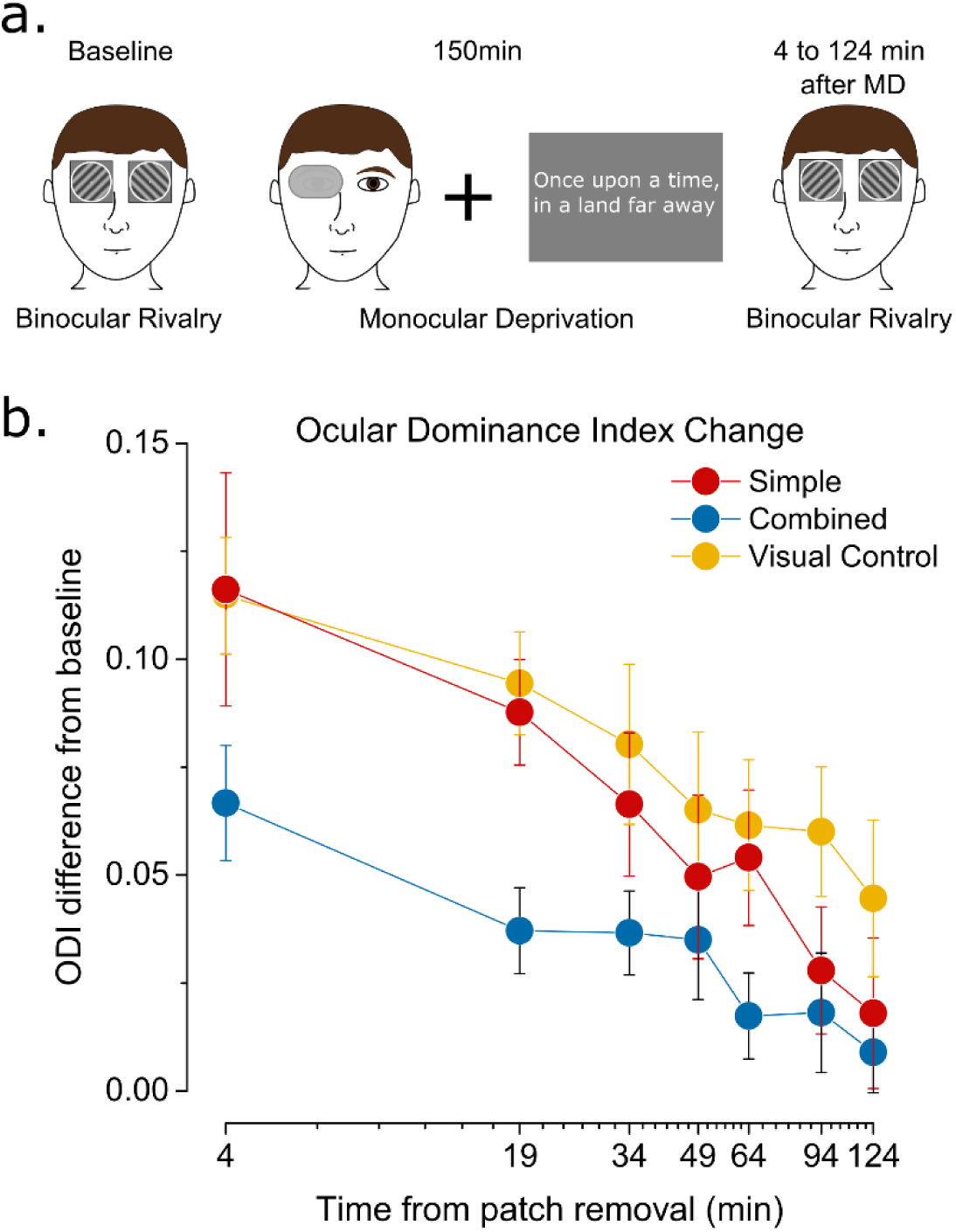
ODI change in the control task. a) In the control study, during the monocular deprivation, 10 participants read short stories on a gray background, displayed a few lines at a time. b) ODI change after monocular deprivation for the simple visual (red symbols), combined (blue symbols) and control (yellow symbols) condition. Error bars represent 1±S.E.M.

Another potential confound in our study was the discrepancy in the working memory and attentional loads between the simple visual task and the combined task during monocular deprivation. Even though there is no clear evidence for a likely impact of working memory or attention on short-term monocular deprivation induced ocular dominance shift, the motor task nevertheless uses resources related to both. To distinguish better whether the effect we see on the ocular dominance shift in the combined condition is due to motor plasticity or merely to attentional engagement, we performed another control experiment in which, during deprivation, the participants performed a working memory task reproducing the demand on working memory and attention as in the simple motor task (Figure 5a). All participants of this attentional control condition also repeated the simple visual condition, and in baseline measurements, the ODI index was strongly correlated for the two conditions (Supplementary Figure 4a, rho = 0.84, p < 0.01), indicating good test-retest reliability. The results show that the ocular dominance shift in this attentional control condition is equivalent to that in the repeated simple visual condition (Figure 5b). A two-way ANOVA (Block x Condition) confirmed that while in both conditions the ocular dominance index changed with blocks (F(5,45) = 14.2, p < 0.001, η^2^p = 0.61), there was no significant effect of condition (F(1,9) = 0.80, p > 0.05, η^2^p = 0.08), nor interaction between the two (F(5,45) = 1.25, p > 0.05, η^2^p = 0.12). Reaction times and accuracy rates in the working memory task confirm that the participants were well engaged with the task (Supplementary Figure 4b & 4c). This suggests that attentional engagement or working memory load does not have an impact on the monocular deprivation effect. In this control experiment, overall, we observed a smaller shift in ocular dominance compared to the main experiment (but no difference between the two conditions). We attribute this difference in the MD effect magnitude to the different experimental setups used for the two experiments, notably a large difference in luminance between the two screens used and a different mirror stereoscope system.

**Figure 5.**
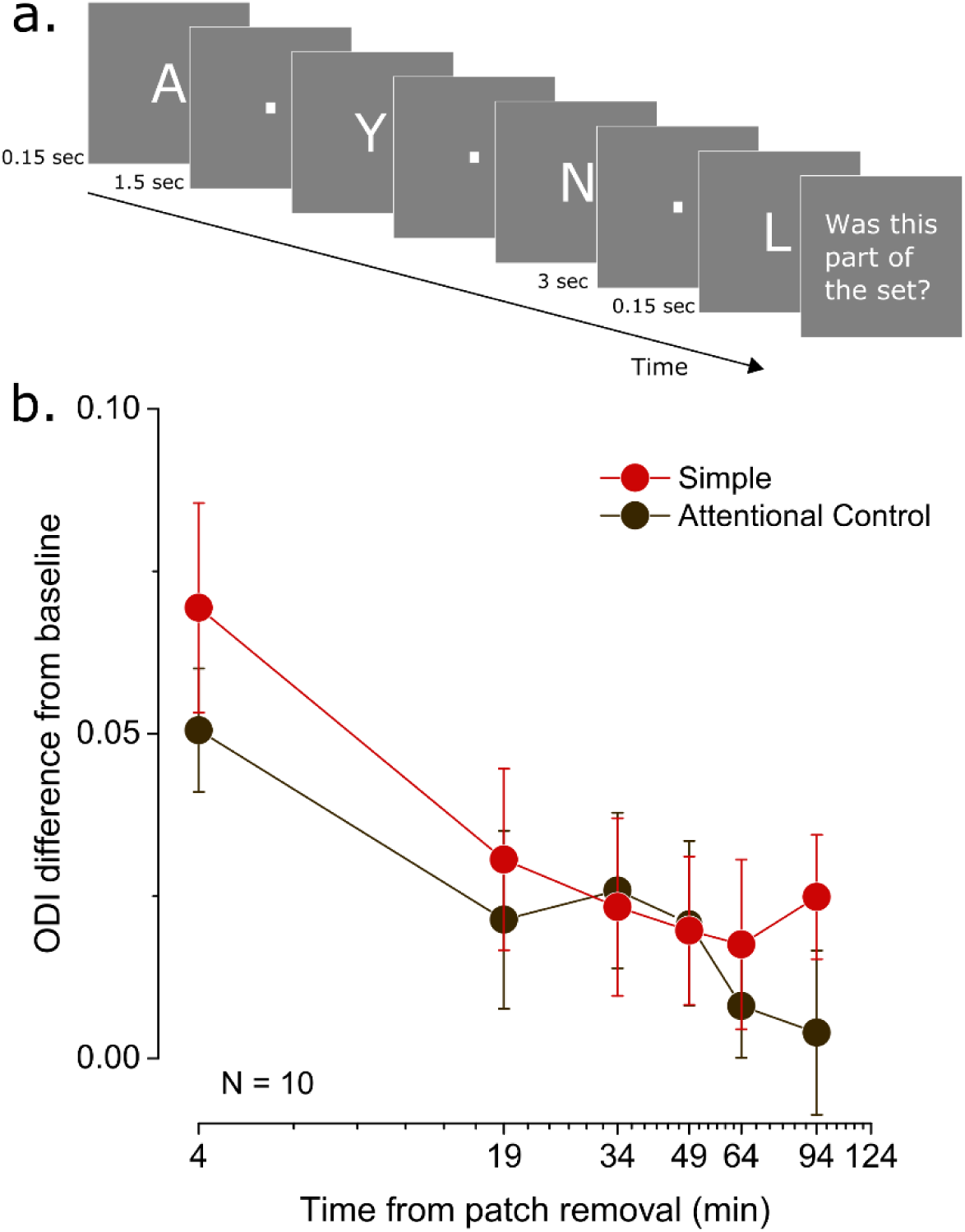
Attentional control task. a) In the attentional control study, 10 participants performed a working memory task during the monocular deprivation. A set of white letters were briefly flashed on a gray background. After a retention period of 3 seconds, participants saw a probe letter flashed and judged whether it was part of the set. They received an auditory feedback after their response. b) ODI change after monocular deprivation for the simple visual (red symbols) and attentional control (brown symbols) condition. Error bars represent 1±S.E.M.

## Discussion

Neuroplasticity in the adult brain has been canonically studied in a modular fashion, supporting the view that plasticity acts locally in different brain areas. Here we investigated for the first time the interaction between two different forms of plasticity by inducing visual and motor plasticity either separately or at the same time in a group of adult human volunteers. We found a unilateral interaction between these two forms of plasticity: when visual and motor plasticity are induced at the same time, visual plasticity is drastically reduced. In two control experiments, we also showed that the reduced visual homeostatic plasticity observed in conjunction with motor plasticity is not attributable to poorer visual stimulation or to the engagement of working memory and attention. Taken together, our results suggest that neuroplasticity is not fully confined to modules in the brain but is subject to interactions, likely to maintain overall homeostasis and prevent excessive instability.

One of the neural mechanisms underlying different forms of plasticity is the balance between excitation and inhibition (E/I), with a predominant role of GABAergic inhibition. Studies from both animal models and humans have shown that decreasing intracortical GABA levels promote visual^41, 42^, motor^24, 25^ and hippocampal^43, 44^ plasticity. Indeed, it has been shown that GABAergic inhibition is also involved in the two paradigms that we used to induce visual and motor plasticity: motor sequence learning induces a decrease of the GABA levels in the motor cortex^26, 27^ and a reduction in visual cortical GABA concentration has been observed after short-term monocular deprivation^14^. It is therefore possible that the observed interaction between motor and visual plasticity is mediated by a disruption of the E/I balance. This hypothesis is consistent with evidence from animal models showing that a localized stroke in S1 or M2 abolishes ocular dominance plasticity^45, 46^. After a stroke, a reduction of GABA concentration occurs at the level of the motor and the sensorimotor cortex. This response enhances plasticity and therefore promotes recovery in the lesioned area^47–50^. We therefore speculate that the negative influence of motor plasticity on visual plasticity that we observed might be due to motor projections on the visual cortex. The disruption in E/I balance in the motor cortex might be transmitted to the visual cortex, disrupting the E/I balance there, in the reverse direction. The reason for this interruptive influence might be maintaining global homeostasis between excitation and inhibition in the brain in order to avoid extreme instability. When it comes to plasticity, the motor cortex might be taking priority and depressing the visual cortex since the visual environment is more stable and motor plasticity is more needed on a general basis. How this short-term interaction is reflected on the consolidation of motor learning, however, is not explored in our study although we see no difference in immediate offline gains.

Positive interactions between the motor and the visual systems have also been observed, one specific case being the influence of locomotion on visual cortical activity and plasticity. Environmental enrichment, with its critical component of physical exercise^51, 52^, promotes the recovery of visual function in adult amblyopic rodents^53^, the restauration of juvenile-like ocular dominance plasticity in healthy adult rodents^54, 55^, and even the preservation of ocular dominance plasticity after a stroke in S1^42, 54^. In humans too, physical exercise has been shown to promote ocular dominance plasticity in both normal sighted^56^ and amblyopic^57^ subjects. One crucial difference between physical activity and motor plasticity (either induced by learning or in response to a stroke) is that during physical activity, the E/I balance changes at the level of the motor cortex are still open to debate^58, 59^. While studies clearly show that motor learning leads to a decrease in GABAergic inhibition in the human motor cortex^24–27^, changes induced by physical activity are not in consensus: although some studies defend a decrease in GABA ^60, 61^ and increased excitability in corticospinal pathway^62–64^, others conclude decreased excitability^65–67^ or higher GABA levels^68^. On the other hand it has been shown that the effect of physical activity on visual plasticity is mediated by decreased GABAergic inhibition in the primary visual cortex^52^ mediated by the activation of a specific disinhibitory circuit (VIP and SST+ interneurons^69, 70^).

While our results show an interaction between motor and visual plasticity, we did not find a correlation between the two. Assuming the plastic capacities of different regions of the brain are linked, we expected to see a positive or negative correlation between the two plasticity scores. The lack of correlation in our results can be due to the different nature of the tasks we used. Motor sequence learning induces a Hebbian form of plasticity that is mainly LTP-like^71–73^. The ocular dominance shift observed after short-term monocular deprivation, on the other hand, is a result of homeostatic mechanisms^12–14, 20^. Homeostatic plasticity is short-lived and aims to preserve a certain E/I balance in the cortex as can be seen with the return to baseline after patch removal. LTP-like plasticity, in motor learning, comes with long-lived changes and goes through a consolidation phase after the initial training. We speculate that, while the lack of correlation between visual and motor plasticity confirms that these two forms of plasticity act at the local level, the interaction between the two indicates the existence of a global homeostatic mechanism regulating local plasticity in separate systems.

Although we included a control for the role of visual stimulation and another for the role of attention and working memory, ruling out a main contribution of these two factors in explaining the observed interaction between visual and motor plasticity, our study lacked a control condition for the role of motor activity. In the combined condition, the participants continuously performed finger taps during the monocular deprivation whereas in the simple visual condition there was no structured motor activity. Despite the fact that the motor sequence learning task we chose does not impose a high level of motor activity on the participants, one might argue that the effect that we see is due simply to motor activity and is not specifically related to motor plasticity. We believe that this is unlikely because of two reasons: first, evidence from previous studies have shown that motor activity has either a boosting^42, 55,56,57^ or a neutral^74^ effect on visual plasticity. Second, in the simple visual condition, during monocular deprivation participants were free to perform motor activity: they all used their phones or computers, performing finger taps analogous to those involved in the motor learning task.

Taken together, our results hint at the existence of a global regulation of plasticity in the brain modulating the local expression of plasticity in separate systems. We speculate that changes in the motor cortex E/I balance mediated by GABA regulation have modulated visual cortex E/I balance in the opposite direction in order to ensure overall stability of activity in the brain and preserve general homeostasis. Given the abundance of environmental conditions requiring motor adaptation in contrast to relative stability of the visual function, we propose that the motor cortex is less restricted in the disruption of E/I balance and hence can take priority over the visual cortex. Further studies focusing on the exact nature of this interaction are in order. Understanding the links between different forms of plasticity is the key for more efficient and innovative approaches to rehabilitative interventions on neurological diseases and brain injury. The ability to tie together motor and visual plasticity may prove useful to recovery from visual or motor dysfunction as seen by the advances in treatment of amblyopia^57^.

## Authors’ contributions

IDS: Designed research, performed research, analyzed data, wrote the paper

CL: Designed research, discussion and interpretation of the results, wrote the paper

## Acknowledgments

This project has received funding from the European Research Council (ERC) under Horizon 2020 research and innovation programme (No 948366 - HOPLA), and from the French National Research Agency (ANR-19-CE28-0008 - PlaStiC and ANR-17-EURE-0017- FrontCog).

## Supplementary Figures

**Supplementary Figure 1.**
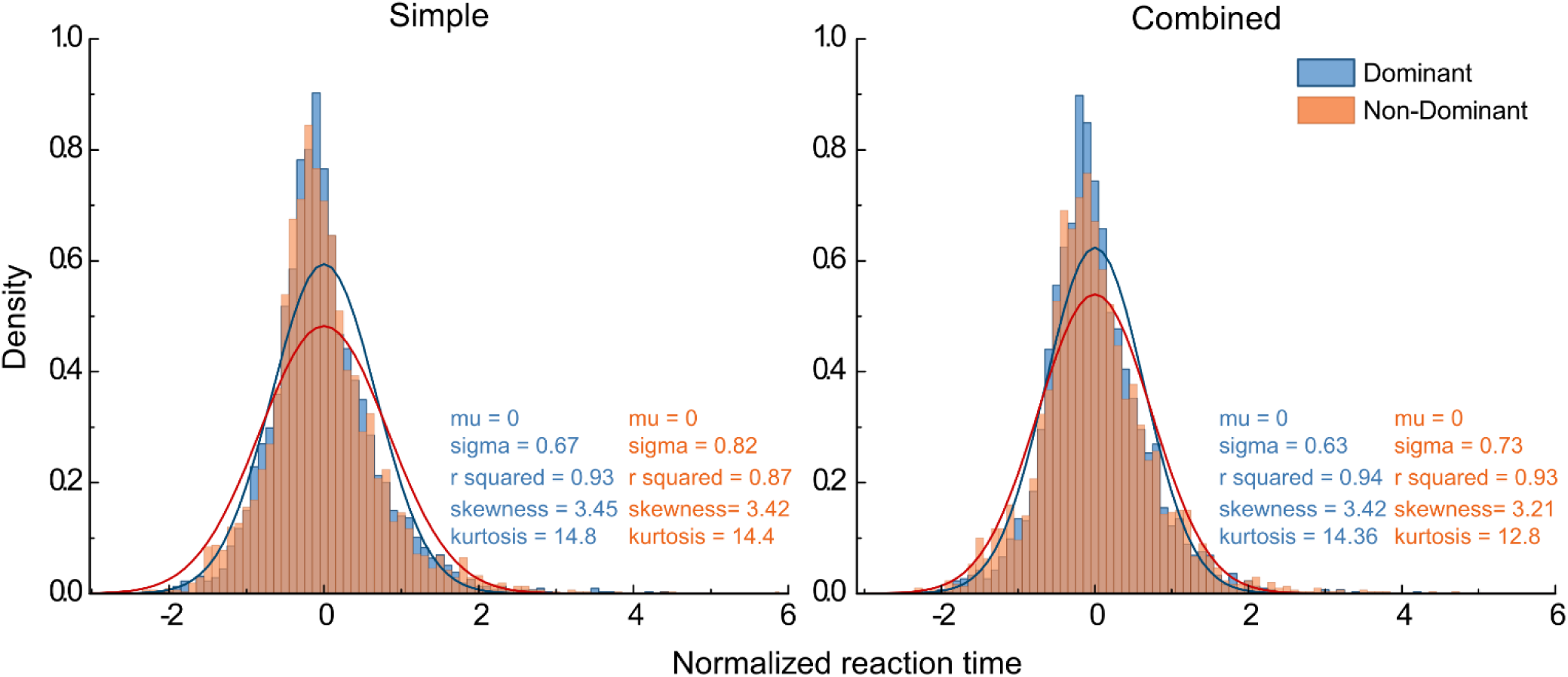
Normalized reaction time distributions. Reaction time distributions, normalized across participants, in the first block of simple and combined motor conditions. The distributions are fitted with a Gaussian function (Eq.3). Skewness and kurtosis are evaluated.

**Supplementary Figure 2.**
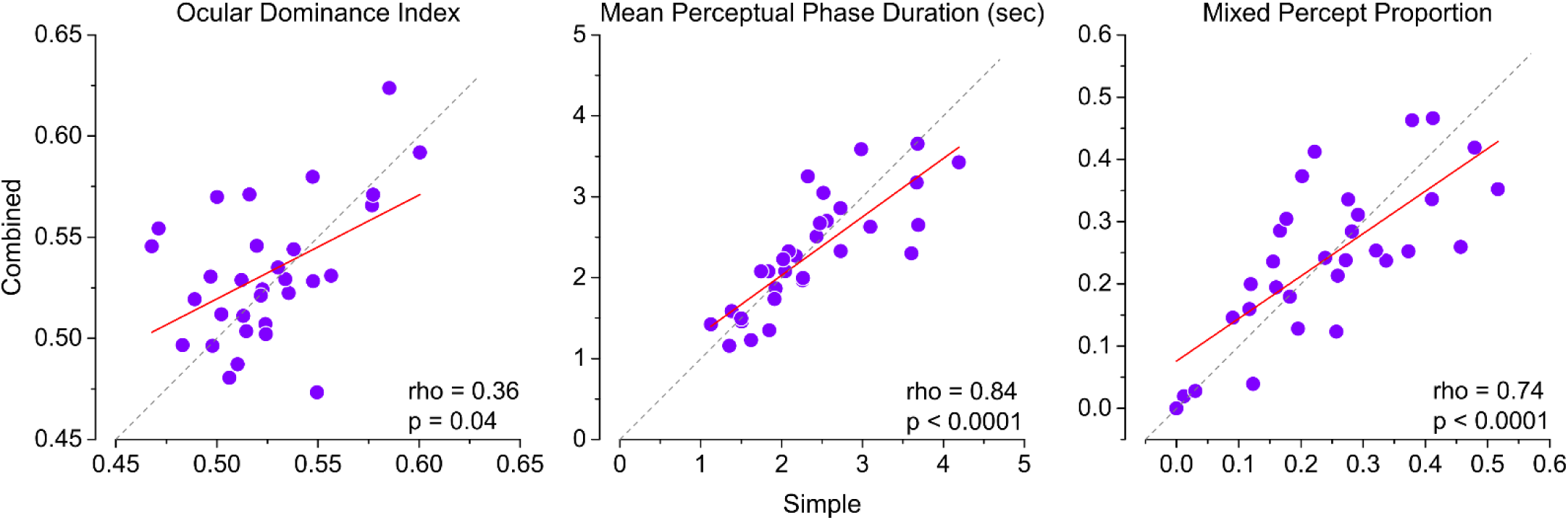
Correlations between the simple and combined conditions for binocular rivalry. a) Correlation between the ODI measured at baseline in the simple vs combined condition (performed on different days). b) Same as A but for mean phase duration of left and right eye perceptions pooled. c) Same as A and B, but for the proportion of mixed percepts.

**Supplementary Figure 3.**
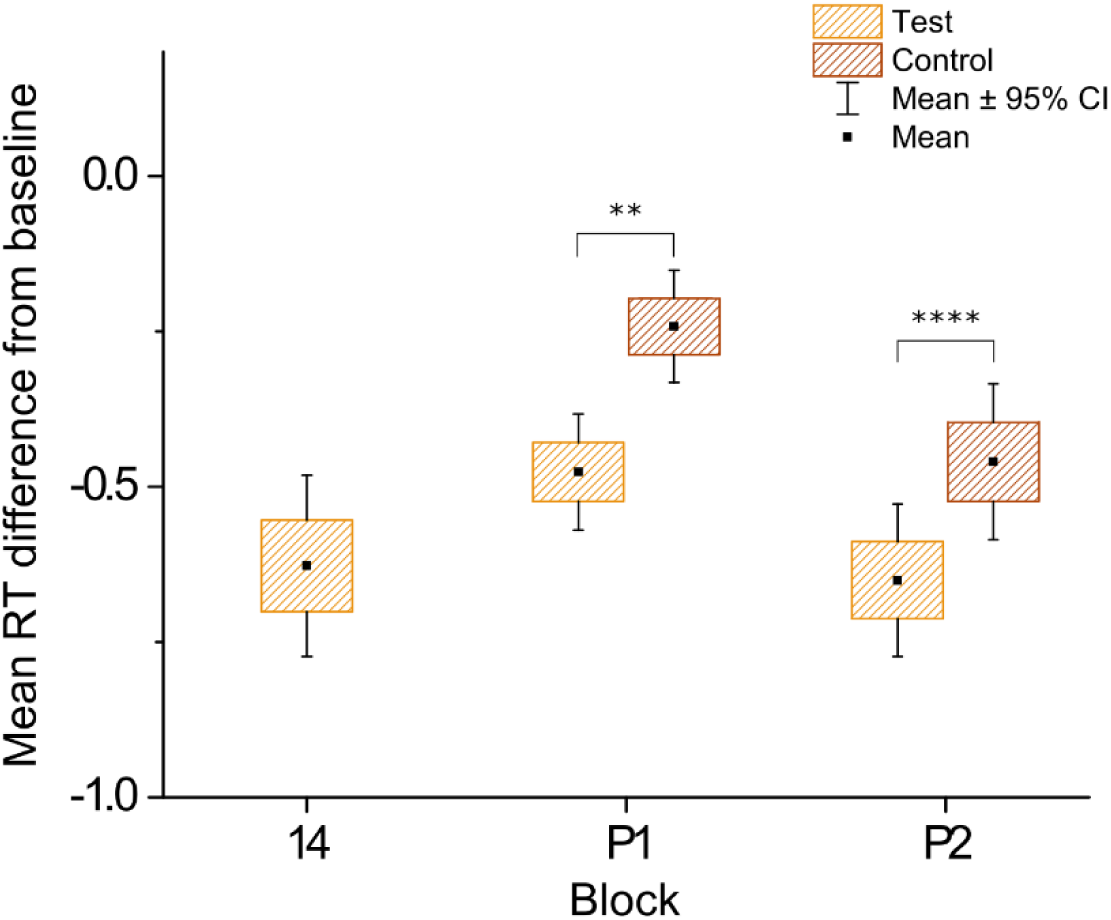
Boxplot of the average reaction time difference from baseline for test and control sequences in simple motor condition. The average reaction time of each participant in the 1^st^ block (test and control sequences pooled) is subtracted from that of 14^th^ block for test sequences, and of P1 (first two trials removed) and P2 blocks, for test and control sequences separately. The black square is the mean. The boxes represent the standard error of the mean. Whiskers denote the 95% confidence intervals around the mean. Significance testing done with Wilcoxon signed rank test. Significance codes: ** p < 0.01, **** p < 0.0001.

**Supplementary Figure 4.**
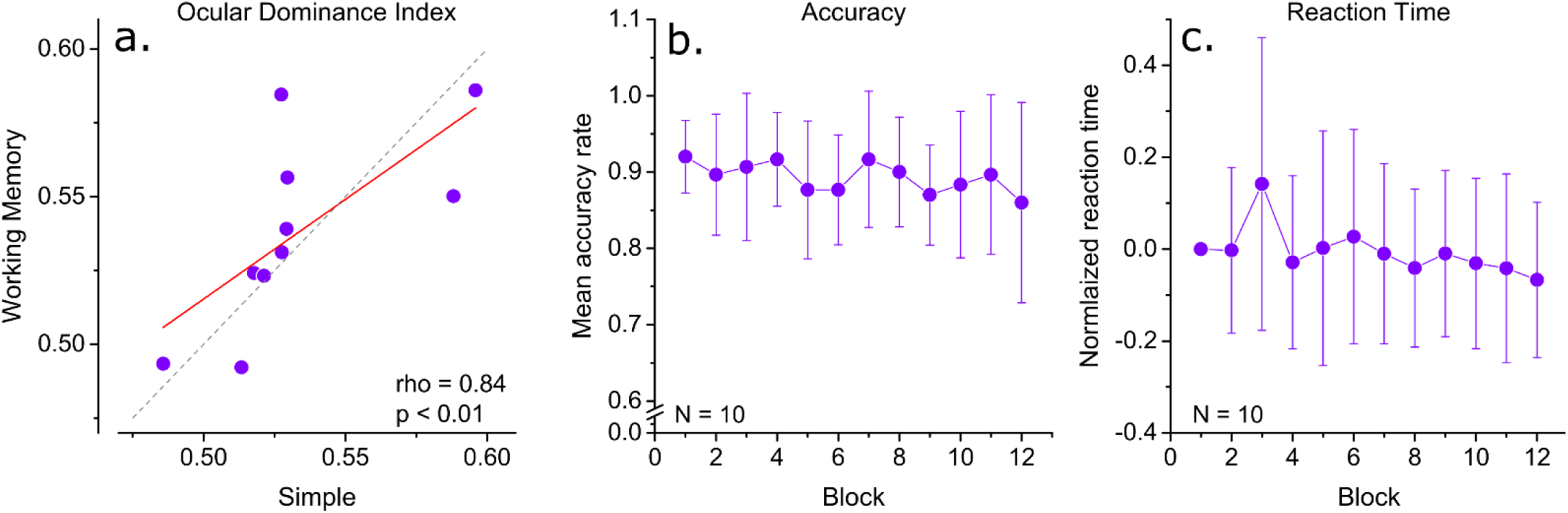
Working memory task. a) Correlation of baseline ocular dominance index in the repeated simple condition and the working memory condition. b) Mean normalized reaction times in working memory task. c) Mean accuracy rate in the working memory task.

